# Sex differences in the human brain transcriptome of cases with schizophrenia

**DOI:** 10.1101/2020.10.05.326405

**Authors:** Gabriel E. Hoffman, Yixuan Ma, Kelsey S. Montgomery, Jaroslav Bendl, Manoj Kumar Jaiswal, Alex Kozlenkov, the CommonMind Consortium, Mette A. Peters, Stella Dracheva, John F. Fullard, Andrew Chess, Bernie Devlin, Solveig K. Sieberts, Panos Roussos

## Abstract

While schizophrenia differs between males and females in age of onset, symptomatology and the course of the disease, the molecular mechanisms underlying these differences remain uncharacterized. In order to address questions about the sex-specific effects of schizophrenia, we performed a large-scale transcriptome analysis of RNA-seq data from 437 controls and 341 cases from two distinct cohorts from the CommonMind Consortium. Analysis across the cohorts identifies a reproducible gene expression signature of schizophrenia that is highly concordant with previous work. Differential expression across sex is reproducible across cohorts and identifies X- and Y-linked genes, as well as those involved in dosage compensation. Intriguingly, the sex expression signature is also enriched for genes involved in neurexin family protein binding and synaptic organization. Differential expression analysis testing a sex-by-diagnosis interaction effect did not identify any genome-wide signature after multiple testing corrections. Gene coexpression network analysis was performed to reduce dimensionality and elucidate interactions among genes. We found enrichment of co-expression modules for sex-by-diagnosis differential expression signatures, which were highly reproducible across the two cohorts and involve a number of diverse pathways, including neural nucleus development, neuron projection morphogenesis, and regulation of neural precursor cell proliferation. Overall, our results indicate that the effect size of sex differences in schizophrenia gene expression signatures is small and underscore the challenge of identifying robust sex-by-diagnosis signatures, which will require future analyses in larger cohorts.

## Introduction

*“The male sex appears in general to suffer somewhat more frequently and to be affected more severely by the dementia praecox” -- Emil Kraepelin, 1919*

Significant sex differences in schizophrenia (called *dementia praecox* at that time) were noted more than 100 years ago by Emil Kraepelin. Since then, multiple epidemiological and clinical studies have described sex differences in age of onset, symptomatology and the course of the disease. First, there is a well established difference in disease incidence among males and females (1). For males, disease onset most commonly occurs in the early twenties. Whereas for females age of onset is bimodal, with an initial mode in the mid-to-late twenties, as well as a second mode in middle age. Second, symptom expression systematically differs between males and females (1). Females are more likely to experience high levels of depressive symptoms, while males are more likely to experience negative symptoms at illness onset. Third, longitudinal studies across 20 years have described sex differences in the presence of psychosis and global outcome (2). Females with schizophrenia are more likely to exhibit fewer psychotic symptoms, as well as better cognitive and global functioning relative to males.

Sex differences in the age of onset, symptomatology and the course of the disease suggest differences in the underlying molecular mechanisms between males and females. Schizophrenia is a multi-factorial neurodevelopmental impairment of the brain that can be attributed to both genetic and environmental factors. Gene expression is a consequence of both the genetic and the environmental factors that contribute to the pathophysiology of the disease. Therefore, transcriptome analysis of the human brain in postmortem studies is a powerful approach for the identification of molecular pathways and signatures associated with schizophrenia. Previous large-scale transcriptome analysis described significant and highly reproducible gene expression changes in schizophrenia (3–5). However, none of these studies explored sex differences that contribute to schizophrenia gene expression dysregulation.

A number of previous studies have performed genome-wide exploration of gene expression in schizophrenia examining sex differences. Qin et al (6) meta-analyzed six microarray datasets from a total from 179 males and 67 females. Gene expression profiling was performed in the dorsolateral or frontopolar prefrontal cortex. They identified significant schizophrenia signatures only in males, while in females, similar analysis did not reveal significant genes after multiple testing corrections. Collado-Torres et al. (7) explored schizophrenia signatures in two brain regions (dorsolateral prefrontal cortex and hippocampus) across 222-238 controls and 132-152 cases (depending on brain region). They found non-significant overlaps between sex and schizophrenia effects for nearly all features. Lack of significant findings might be due to limited power, indicating the need to examine sex differences in larger cohorts.

To address this knowledge gap and increase statistical power, in this study we performed a large-scale transcriptome analysis of sex differences in schizophrenia using 437 controls and 341 cases from the CommonMind Consortium RNA-seq collection (3, 8). We specifically address the following questions: What are the sex differences in gene expression in the brain? Are there schizophrenia genes that are affected differently in males compared to females? And, if so, do those differences affect specific molecular pathways and co-expression modules?

## Methods and Materials

### Description of cohorts

Experimental methods for generating the CommonMind Consortium RNA-seq dataset from the dorsolateral prefrontal cortex are described in release v2 (8). The collection involves two cohorts derived from four brain banks: ***(A)*** The initial cohort comprised of samples from the Mount Sinai School of Medicine Brain Bank, University of Pennsylvania Brain Bank and University of Pittsburgh Brain Bank (MSSM-Penn-Pitt); and ***(B)*** the cohort ascertained from the National Institute of Mental Health’s Human Brain Collection Core (NIMH-HBCC). See https://www.synapse.org//#!Synapse:syn2759792/wiki/194729 for further details. In this study, we overall included 281 females and 497 males samples (**Supplementary Table 1**).

### Processing RNA-seq data

RNA-seq data were generated as previously described (8) and were processed as follows. The raw reads were trimmed with Trimmomatic (v0.36) (9) and then mapped to human reference genome GRCh38.v24 (ftp://ftp.ebi.ac.uk/pub/databases/gencode/Gencode_human/release_24/GRCh38.primary_assembly.genome.fa.gz) using STAR (v2.7.2a) (10). The BAM files that were generated contain the mapped paired-end reads, including those spanning splice junctions. Following read alignment, expression quantification was performed at the gene level using featureCounts (v1.6.3) (11). Gene quantifications correspond to GENCODE v30 (ftp://ftp.ebi.ac.uk/pub/databases/gencode/Gencode_human/release_30/gencode.v30.annotation.gtf.gz). Quality control metrics were reported with Picard (v2.20.0). The full RNA-seq pipeline is implemented in Nextflow (12) and is available at https://github.com/CommonMindConsortium/RAPiD-nf. Analysis used log_2_ counts per million (CPM) following TMM normalization (13) implemented in edgeR (v3.22.5) (14). Genes with over 0.5 CPM in at least 30% of the samples in both cohorts were retained.

### Computational deconvolution

Dtangle (15) was used to estimate the cell type composition in the bulk RNA-seq data using a reference panel composed of four cell components generated based on fluorescence activated nuclear sorting (FANS). The reference panel included GABAergic neurons (GABA), glutamatergic neurons (GLU), oligodendrocytes (Olig), and the remaining fraction consists of mostly microglia and astrocytes (MgAs). Data were generated from the dorsolateral prefrontal cortex of a subset of 32 MSSM samples. The raw reads were preprocessed using the pipeline described above. The mean log_2_ CPM for each gene for each cell type was used as the reference panel for deconvolution.

### Generation of FANS reference panel

Individual, 50mg aliquots, of frozen brain tissue were homogenized in cold lysis buffer (0.32M Sucrose, 5 mM CaCl2, 3 mM Magnesium acetate, 0.1 mM, EDTA, 10mM Tris-HCl, pH8, 1 mM DTT, 0.1% Triton X-100). Samples were filtered through a 40μm cell strainer and underlaid with sucrose solution (1.8 M Sucrose, 3 mM Magnesium acetate, 1 mM DTT, 10 mM Tris-HCl, pH8) prior to ultracentrifugation at 107,000 g for 1 hour at 4°C in a swing bucket rotor. Pellets were resuspended in 500μl DPBS (supplemented with 0.1% BSA) and incubated with anti-NeuN (1:1000, PE conjugated, Millipore Cat FCMAB317PE), anti-SOX6 (16) and anti-SOX10 (17) antibodies for 1 hour at 4°C with end-over-end rotation, in the dark. Following incubation in primary antibodies, samples were subjected to a second ultracentrifugation step prior to incubation in secondary antibodies, as above (18).

Immediately prior to FANS sorting, DAPI (Thermoscientific) was added to a final concentration of 1μg/ml. GABAergic neurons (DAPI+ NeuN+ SOX6+), Glutamatergic neurons (DAPI+ NeuN+ SOX6-), oligodendrocytes (DAPI+ NeuN-SOX10+) and microglia/astrocytes (DAPI+ NeuN-SOX10-) nuclei were sorted into individual tubes using a FACSAria flow cytometer (BD Biosciences).

FANS sorted nuclei for RNA-seq were collected in PicoPure (Applied Biosystems) extraction buffer and were incubated at 42°C for 30 min under shaking at 850 rpm, before storage at −80°C. RNA extraction was performed using the PicoPure RNA Isolation kit (Applied Biosystems) and RNA-seq libraries generated using the SMARTer cDNA synthesis kit (Takara), according to manufacturer’s instructions. Libraries were sequenced at New York Genome Center on the NovaSeq platform (Illumina) obtaining 100bp paired-end reads.

### Covariate exploration

Observed gene expression measurements from RNA-seq can be affected by biological and technical factors. To identify important covariates, we examined the correlation between multiple variables and the gene expression data, as well as the correlation between multiple biological and technical variables. While evaluating the correlation between two continuous variables is trivial, evaluating the correlation between two categorical variables, or including a variable with multiple dimensions (i.e. cell type fraction) is more complicated. We applied a generalization of the standard correlation by using canonical correlation analysis and reporting Cramér’s V statistic (19), which is the fraction of the maximum possible correlation. When comparing two continuous variables, this gives the same value as the typical correlation.

The fraction of variance in cell type composition explained by other variables was evaluated with the variancePartition package (20). All variables were modeled as fixed effects.

### Differential expression

Analysis was performed using dream (21) built on top of limma-voom (22). In addition to diagnosis and sex, the following variables were included as covariates: RIN, intronic rate, intragenic rate, intergenic rate, rRNA rate, Institution, age of death and cell type composition. These covariates were identified empirically using the Bayesian Information Criterion (23) to identify important variables in each cohort separately. The union of the variables identified in either cohort was then used in the analysis.

Since age of death and cell type composition varied across institutions, interaction terms between institution and age of death and intuition and cell type composition were used. While computational deconvolution was used to estimate the fraction of four cell types, the constraint that the four fractions sum to 1.0 for each sample means that the fractions really span only three dimensions (i.e., knowing three determines the fourth). Including this set of low rank covariates in a regression model is problematic. To address this issue, the four fractions were transformed into three variables using the isometric log ratio (24) to create covariates `cellFrac_ilr_1, cellFrac_ilr_2, cellFrac_ilr_3`. This transformation is invariant to scaling and reordering of the cell fraction variables; however, each resulting variable is a function of multiple cell fractions so they can’t be interpreted individually.

The regression formula used for differential expression analysis was:

~~~
     ~ Diagnosis + Sex + RIN + IntronicRate + IntragenicRate +
     IntergenicRate + rRNARate + Institution*(ageOfDeath + cellFrac_ilr_1 +
     cellFrac_ilr_2 + cellFrac_ilr_3)
~~~

in which RIN is the RNA integrity number measured from the physical RNA, intronic rate is the fraction of reads mapping to introns, intragenic rate is the fraction of reads mapping to exons or introns, intergenic rate is the fraction of reads mapping to intergenic regions, rRNA rate is the fraction of reads mapping to ribosomal RNA. With the exception of RIN, these other metrics were computed from RNA-seq reads by Picard.

The sex-by-diagnosis interaction analysis used the terms `Diagnosis + Sex + Diagnosis:Sex` and tested the `Diagnosis:Sex` while accounting for the covariates above.

Results from the MSSM-Penn-Pitt and NIMH-HBCC cohorts were combined with a fixed effects meta-analysis using the regression coefficients from each cohort and their respective standard errors with the metafor package (25). The fraction of replicated differentially expressed genes was estimated using the 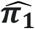 statistic from the qvalue package (26).

### Gene set analysis

Testing gene sets for enrichment of differential expression signatures was performed with zenith (https://github.com/GabrielHoffman/zenith), which is a slight modification of camera (27) to be compatible with dream (21). It is a competitive gene set test that uses the full spectrum of t-statistics from the differential expression analysis without specifying a p-value or false discovery rate (FDR) cutoff to call a gene differentially expressed. This approach explicitly models the empirical correlation between genes in each gene set to control the false positive rate accurately (27). When performing gene set analysis on the differential expression signature from the metaanalysis of the two cohorts, modelling this correlation structure while controlling the false positive rate is challenging due to the two datasets. Instead, gene set analysis was performed on each cohort separately and results for each gene set were combined using a fixed effect meta-analysis in the metafor package (25). Gene sets from Gene Ontology (28) were obtained using EnrichmentBrowser (29).

### Similarity between differential expression signatures

To evaluate the robustness of the schizophrenia differential expression signature, the signature was compared to signatures from other datasets. Differential expression signatures measuring differences between individuals with schizophrenia to controls from the current data and were compared to ***(1)*** schizophrenia signatures from the CommonMind Consortium v1 release (3); ***(2)*** schizophrenia, bipolar disorder and autism spectrum disorder from the PsychENCODE Consortium (4); and ***(3)*** Alzheimer’s disease signatures from the AD Knowledge Portal (30). The Alzheimer’s disease signature was included as a negative control because it is a disease of the brain but acts via a very different molecular etiology. All differential expression signatures were from RNA-seq data, except the NIMH-HBCC cohort from the CommonMind Consortium v1 release, which was from Illumina HumanHT-12_V4 Beadchip microarrays (3).

Similarity between a pair signatures was computed as the Spearman correlation between the t-statistics. Hierarchical clustering was performed with the `“ward.D2”` method. Bootstrap resampling using pvclust (31) was used to evaluate the stability of the clusters. All clusters had approximately unbiased probability of ≥ 0.85, indicating high stability.

### Network analysis

Multiscale embedded-gene coexpression network analysis (MEGENA) (32) was performed on expression residuals to identify modules of highly co-expressed genes in MSSM-Penn-Pitt and NIMH-HBCC cohorts, respectively. Expression residuals were produced by regressing out covariates using the formula described in the *“Differential expression”* section and adding back terms corresponding to the intercept, diagnosis, sex and diagnosis-by-sex interaction. Gene-gene similarities were measured by Pearson correlation. Then gene pairs were selected with significant correlations based on a cutoff of 0.05 after FDR correction. These gene pairs were embedded onto a topological sphere to construct a planar filtered network. Multiscale clustering analysis was performed on the planar filtered network to determine the module compactness, contributing to a hierarchy of parent and child modules. The key driver genes of each module were identified by multiscale hub analysis using Fisher’s inverse Chi-square approach in MEGENA.

We explored the overlap of each module with multiple gene sets and external data as described below. Significant associations were determined based on an FDR cutoff of 5% in each analysis. For each module, the following analyses were performed:

- <u>Overlap with differential expression signatures</u>: The differential expression signatures from sex, diagnosis, and sex-by-diagnosis interaction were tested for enrichment in each module using cameraPR in the limma package (33).
- <u>Overlap with module genes</u>: Overlapping modules among the MSSM-Penn-Pitt and NIMH-HBCC networks were determined based on Fisher’s exact test.
- <u>Gene set enrichment analysis</u>: The modules were functionally annotated by gene set enrichment of biological pathways from MSigDB (34) based on Fisher’s exact test.
- <u>Overlap with risk variation</u>: MAGMA gene-set analysis (35) was performed for the PGC2 schizophrenia associations (36).

## Results

### Cell type composition and confounding

While the biological variables of interest are sex and diagnosis, identifying and accounting for confounding variables is essential to reduce spurious correlations. Thus, it is important to understand which experimental and biological variables are associated with sex in our dataset to correct for them. Sex is correlated with multiple variables when combining individuals across the MSSM-Penn-Pitt and NIMH-HBCC cohorts (**Figure 1A**). Females tend to be older, are underrepresented in Pitt and NIMB-HBCC brain banks (at 26.5% and 28.1% respectively) and have a lower estimated fraction of GABAergic neurons (**Supplementary Figures 1,2**). Diagnosis shows lower correlation with these variables than sex.

**Figure 1.**
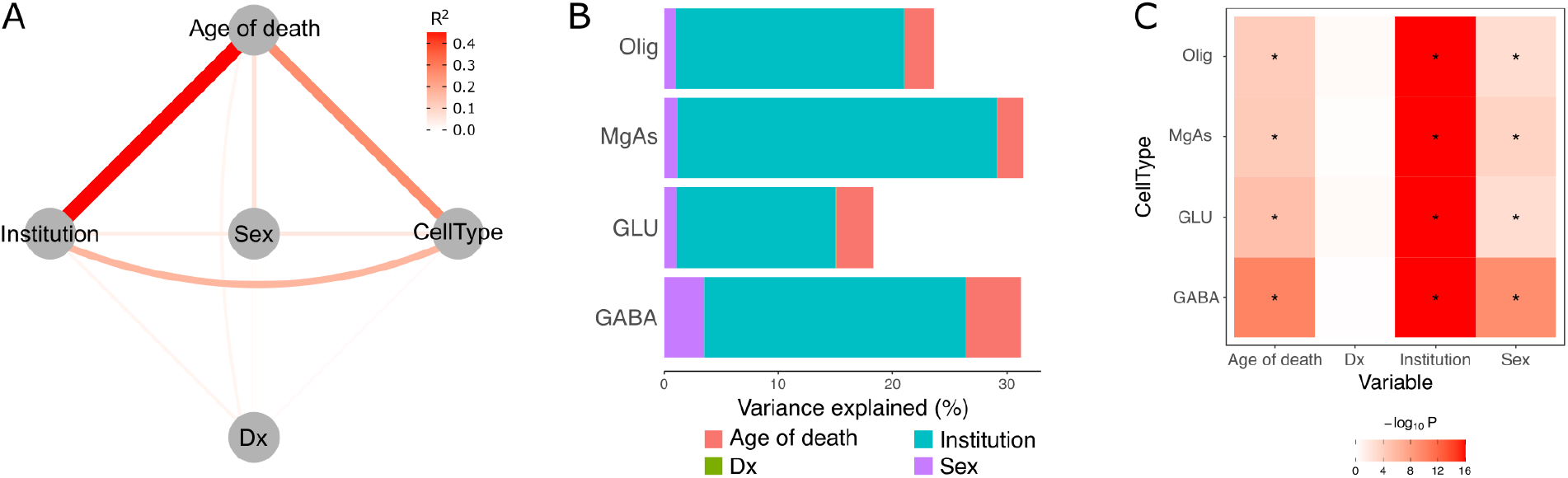
Association with estimated cell type composition. **A)** Network showing squared correlation between each pair of variables. Color and thickness of the line corresponds to the magnitude of the squared correlation. **B)** Variance partitioning analysis shows the fraction of variance in each cell type component explained by age of death, diagnosis, institution and sex. The remaining variation is the residual variance not explained by these variables. **C)** Hypothesis testing using ANOVA identified variables explaining a significant fraction of variance in each cell type component. Stars indicate tests passing the 5% Bonferroni cutoff for 16 tests.

Another way of examining the correlation between these variables is by evaluating the fraction of variation in cell type composition explained by age of death, diagnosis, institution (i.e. brain bank) and sex (**Figure 1B**). Across all 4 cell types in the reference panel (GABA, Glu, Olig and MgAs), institution explained the most of the variation in cell type composition, followed by age of death and sex. Each of these variables explained a significant fraction of the variation in cell type composition even after Bonferroni correction (**Figure 1C**). Notably, diagnosis explained very little variance in cell type composition, and its contribution was not statistically significant. Based on this analysis, to evaluate sex differences in schizophrenia and to avoid spurious associations driven by confounds, we included age of death, institution, and cell type composition as covariates in the statistical model.

### Transcriptomic signature of schizophrenia

Differential expression analysis between schizophrenia and controls identified 217 genes in the MSSM-Penn-Pitt cohort and 1,706 genes in the NIMH-HBCC cohort at FDR 5%. Despite the substantial difference in the number of genome-wide significant differentially expressed genes (DEGs), the disease signatures were remarkably concordant (**Figure 2A**). The t-statistics from the two cohorts had a large Spearman correlation of 0.343 (p < 1 x 10^−300^). Moreover, considering MSSM-Penn-Pitt as the discovery cohort, the replication rate in NIMH-HBCC estimated using the π_1_ statistic was 89.7%. Considering NIMH-HBCC as the discovery cohort, the replication rate in MSSM-Penn-Pitt was estimated to be 46.1%. Combining results from the two cohorts using a fixed effects meta-analysis identified 2,209 significant differentially expressed genes at FDR 5% (**Figure 2B**).

**Figure 2.**
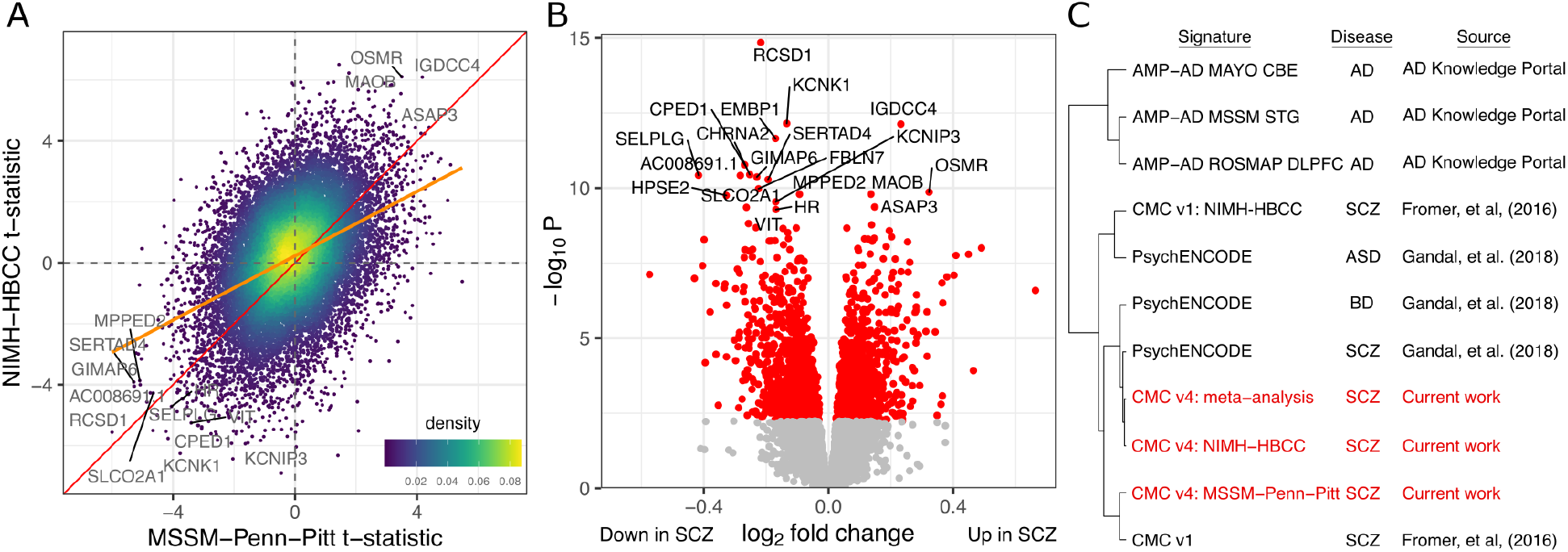
Differential expression between schizophrenia and controls. **A)** Concordance analysis showing t-statistics for each gene from MSSM-Penn-Pitt and NIMH-HBCC cohorts. Orange line indicates best fit from linear regression. Color of points indicates the density in the local region. **B)** Volcano plot of meta-analysis combining both cohorts. Red points indicate FDR < 0.05. **C)** Clustering of differential expression signatures from the current work compared to previously published disease signatures from post mortem brain including schizophrenia (SCZ), bipolar disorder (BD), Alzheimer’s disease (AD) and autism spectrum disorder (ASD). Alzheimer’s’ signatures are from the cerebrum (CBE), superior temporal gyrus (STG) and dorsolateral prefrontal cortex (DLPFC). All other signatures are from the DLPFC.

The schizophrenia signatures from the two cohorts and the combined meta-analysis were concordant with existing schizophrenia signatures from the CommonMind Consortium (3) and PsychENCODE (4), and to a lesser degree with bipolar disorder and autism spectrum disorder signatures from PsychENCODE (4) (**Figure 2C**, **Supplementary Figure 3**). Importantly, there was little concordance with 3 disease signatures from Alzheimers’ disease (**Figure 2C**) which, despite being a disease of the brain, is known to have a distinct molecular etiology.

### Differential expression between males and females

Differential expression analysis between males and females identified 482 genes, including 418 autosomal genes, in the MSSM-Penn-Pitt cohort and 148 genes, including 98 autosomal genes, in the NIMH-HBCC cohort at FDR 5%. Similar to the disease analysis, the sex signatures were notably concordant and had a Spearman correlation between t-statistics of 0.15 (p < 1.23 x 10^−96^) for all genes and 0.14 (p < 1.6 x 10^−85^) for autosomal genes (**Figure 3A**). Considering MSSM-Penn-Pitt as the discovery cohort, the estimated replication rate in NIMH-HBCC was 47.5% (39.5% for autosomal genes). Considering NIMH-HBCC as the discovery cohort, the replication rate in MSSM-Penn-Pitt is estimated to be 52.8% (28.4% for autosomal genes). Notably, differential expression identified on sex chromosomes (chromosomes X and Y) in one cohort was more likely to replicate. Combining results from the two cohorts using a fixed effects meta-analysis identified 686 significant DEGs at FDR 5% (**Figure 3B**), including 606 autosomal genes (**Figure 3C**).

**Figure 3.**
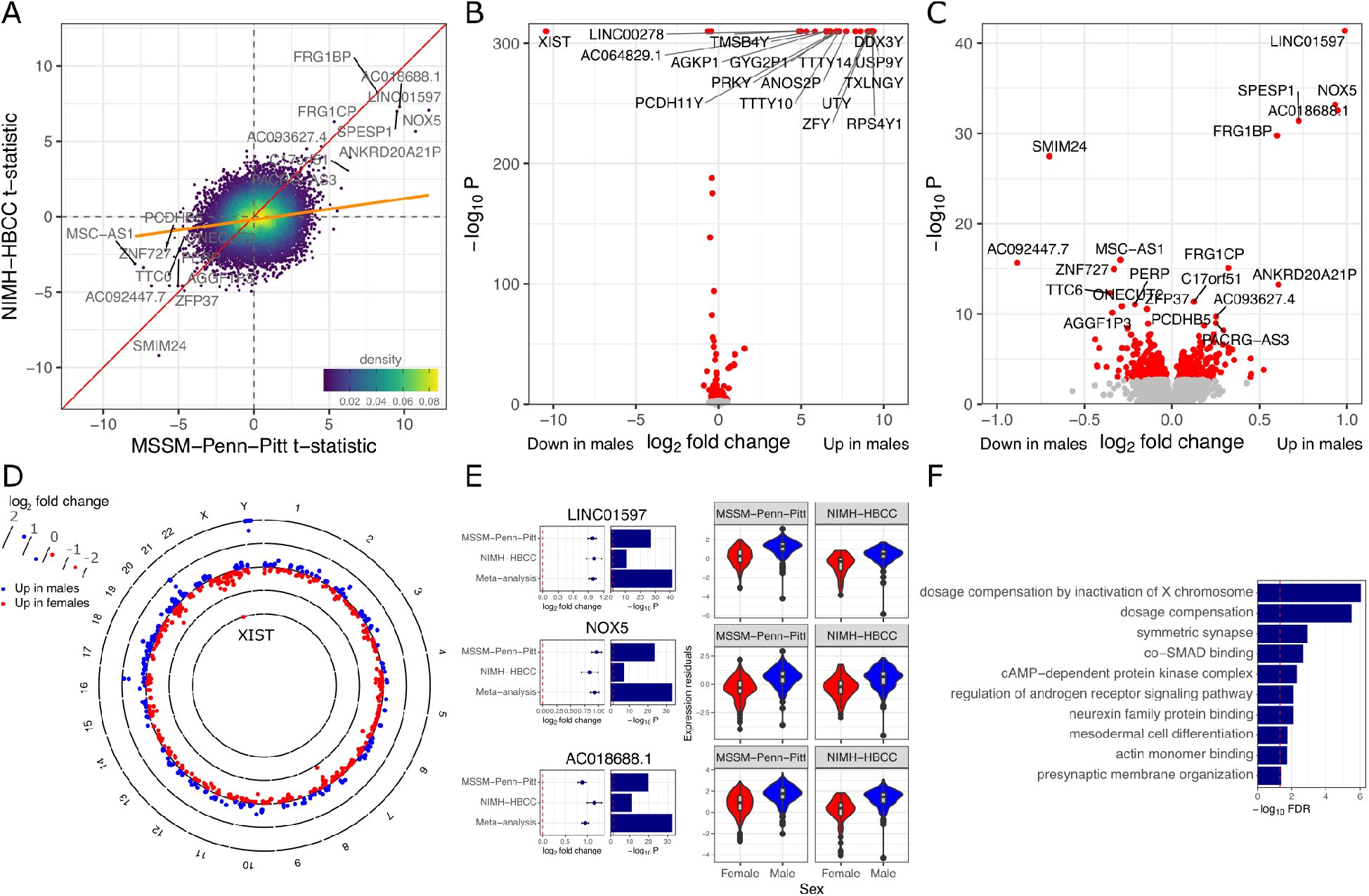
Differential expression between males and females. **A)** Concordance analysis showing t-statistics for each gene from MSSM-Penn-Pitt and NIMH-HBCC cohorts. The orange line indicates best fit from linear regression. Color of points indicates the density in the local region. **B)** Volcano plot of meta-analysis combining both cohorts for all genes. Red points indicate FDR < 0.05. **C)** Volcano plot showing only autosomal genes. **D)** Circos plot of log_2_ fold changes by chromosome position. Only genes with FDR < 5% are shown. Log_2_ fold changes are thresholded to be between −2 and 2, but this only affects genes on the sex chromosomes. **E)** Examples of differentially expressed autosomal genes. Left panel shows log_2_ fold changes in each cohort and meta-analysis with error bar indicator ± 1 standard error unit and corresponding −log_10_ p-value. Right panel shows violin plot of expression, after regressing out covariates, stratified by cohort and sex. **F)** Gene set analysis identifies Gene Ontology annotations enriched for genes differentially expressed between sexes.

As expected, the effect sizes were substantially larger for DEGs on the sex chromosomes compared to autosomes (**Figure 3D**). The top autosomal DEGs identified by meta-analysis showed very consistent effects sizes across cohorts (**Figure 3E**). Unsurprisingly, gene set enrichment analysis identified molecular pathways involved in dosage compensation and androgen signaling (**Figure 3F**). Intriguingly, sex differences in the brain transcriptome identified genes that affect neurexin family protein binding and synaptic organization.

### Effect of sex on disease differential expression signature

We then examined whether the effect of disease differs between males and females by statistically testing an interaction term between sex and diagnosis. In each cohort, no genes passed an FDR threshold of 10%. The t-statistics from the two cohorts showed a low, yet significant, level of similarity (Spearman correlation = 0.054, p < 5.3 x 10^−14^, **Figure 4A**). We also performed differential expression analysis for diagnosis, separately by sex. The cross-cohort meta-analysis shows high concordance between the schizophrenia signature in males and females with a Spearman correlation of 0.453 (p < 1 x 10^−300^); no significant difference in effect sizes was identified (**Supplementary Figure 4**). Combining results from the two cohorts in a meta-analysis yielded no genes passing an FDR threshold of 5% (**Figure 4B**); only ALKBH3 passes a 10% FDR cutoff. The effect size of −0.179 for ALKBH3 indicated that the schizophrenia-vs-control effect size in males was less than the effect size in females (**Figure 4C**). The low concordance across cohorts and the lack of any significant finding at FDR 5% (and only one significant gene at FDR 10%) indicates a limited power to identify sex differences in the schizophrenia signature in the current data.

**Figure 4.**
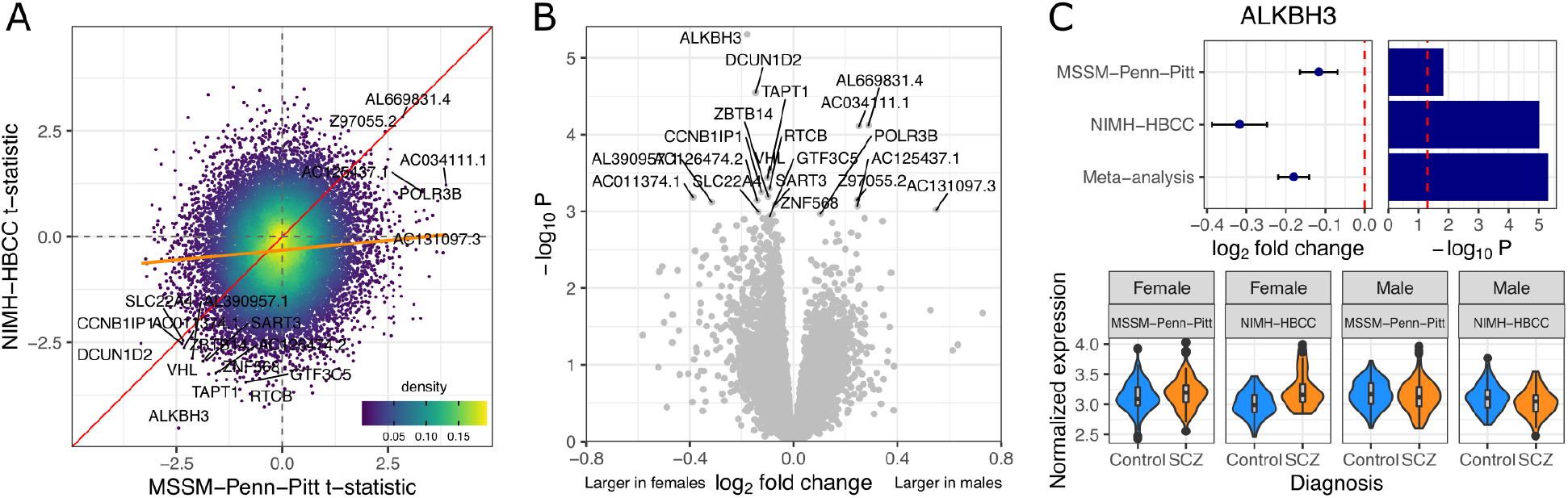
Interaction analysis for effect of sex on schizophrenia signature. **A)** Concordance analysis showing t-statistics for each gene from MSSM-Penn-Pitt and NIMH-HBCC cohorts. Orange line indicates best fit from linear regression. Color of points indicates the density in the local region. **B)** Volcano plot of meta-analysis combining both cohorts for all genes. **C)** Top panel shows results for the top gene, ALKBH3, showing log_2_ fold change and p-values for each cohort and meta-analysis. Bottom panel shows violin plots of ALKBH3 expression residuals stratified by sex, cohort and diagnosis.

### Network analysis identifies gene modules

To further understand how sex differences contribute to schizophrenia gene expression, we used multiscale embedded-gene coexpression network analysis (MEGENA) to identify co-expressed gene modules and then characterize their enrichment for differential expression signatures for diagnosis, sex, and sex-by-diagnosis interaction. A total of 1,226 and 1,396 hierarchical (i.e. parent-child) modules were identified from the MSSM-Penn-Pitt and NIMH-HBCC cohorts, respectively. These modules were then ranked by their association with diagnosis, sex, and sex-by-diagnosis signatures by performing the enrichment analysis with the respective differential expression signature. A total of 253 significantly enriched modules were identified in the MSSM-Penn-Pitt cohort at FDR 5% (212 for diagnosis, 7 for sex, and 208 for sex-by-diagnosis) (**Figure 5A**). Similar analysis in the NIMH-HBCC cohort identified 236 significantly enriched modules at FDR 5% (184 for diagnosis, 8 for sex, and 201 for sex-by-diagnosis) (**Figure 5B**).

**Figure 5.**
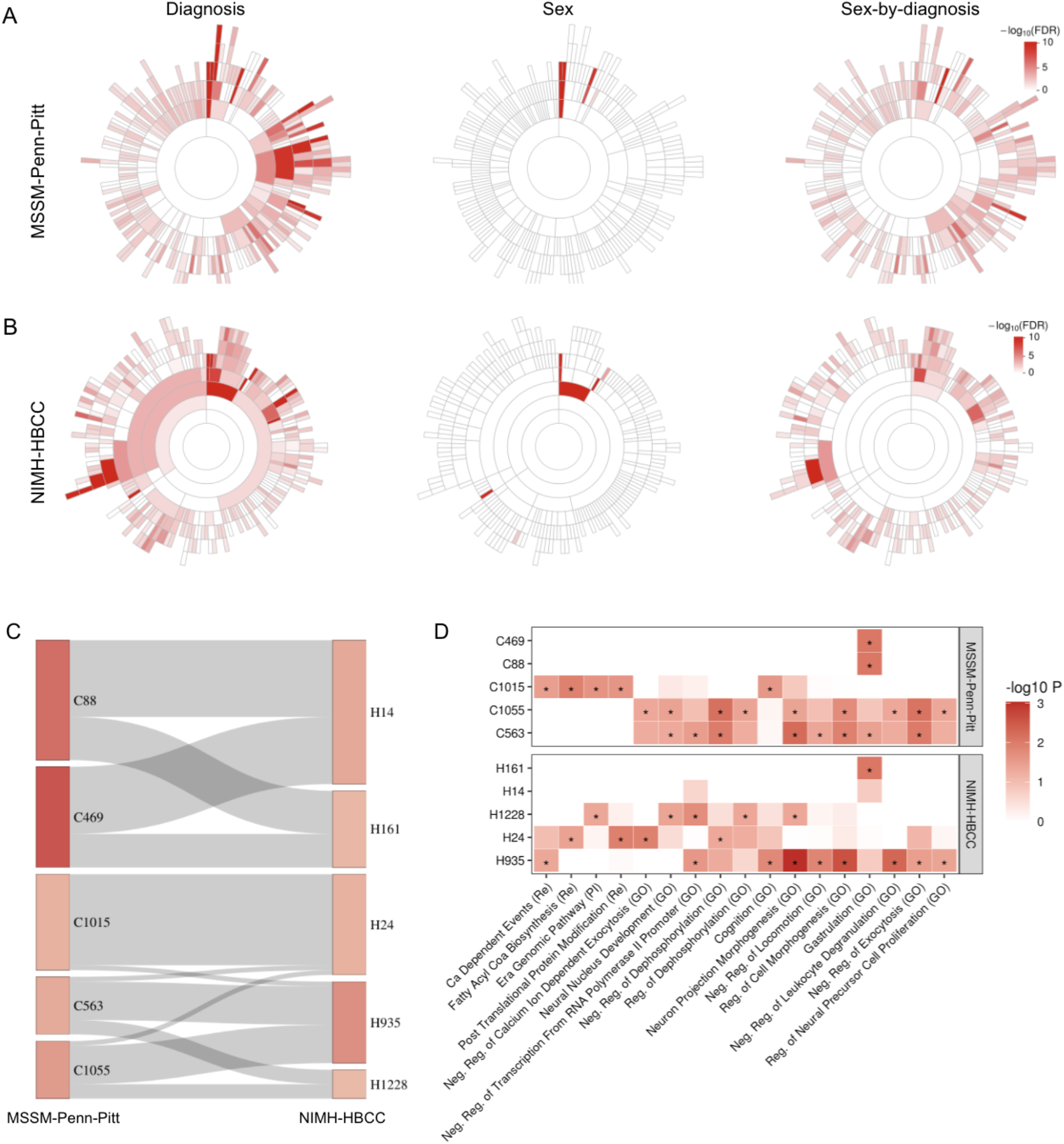
Gene co-expression modules associated with sex and schizophrenia in. **A)** MSSM-Penn-Pitt and **B)** NIMH-HBCC. Enrichment of the differential expression signatures of diagnosis (schizophrenia; left panel), sex (middle panel), and sex-by-diagnosis (interaction; right panel) in all of the significantly enriched modules in MSSM-Penn-Pitt and NIMH-HBCC cohorts, respectively. Sunburst plot showing the hierarchical structure of enriched modules. Each sector represents module enriched with either diagnosis, sex, or sex-by-disease. The most inside sector represents the root module. The inside sectors represent parent modules, and the outside sectors represent child modules. Color intensity is proportional to −log_10_ FDR for enrichment of each module with differential expression signatures. **C)** Sankey plot showing a high overlap among the top 5 interaction enriched modules from the MSSM-Penn-Pitt and NIMH-HBCC cohorts. Color intensity is proportional to −log_10_ FDR from the interaction enrichment analysis (same as the color scale from **(A)** at the right panel. The width of the ribbon represents the number of overlapping genes between two correlated modules. **D)** Gene set enrichment analysis for the top 5 interaction enriched modules for both cohorts (shown in **(C)**). Color intensity is proportional to −log_10_ p-value and asterisk shows enrichments that are significant at nominal P < 0.05.

Sex-by-diagnosis interaction associated modules were more likely to be enriched for diagnosis differential expression signatures for both MSSM-Penn-Pitt and NIMH-HBCC cohorts (Spearman rho = 0.68 and 0.60 at P < 2.2 x 10^−16^, respectively) (**Supplementary Figure 5**). Only a small number of modules were enriched for sex signatures (**Figure 5A, B** middle panel), which show only partial overlap with the diagnosis effects. We checked the replication of sex-by-diagnosis interaction associated modules across the two cohorts. Considering interaction enriched modules of the MSSM-Penn-Pitt cohort as the reference, the replication rate in NIMH-HBCC (based on π_1_ statistic) was 39%. Considering interaction enriched modules of the NIMH-HBCC cohort as the reference, the replication rate in MSSM-Penn-Pitt was estimated to be 41%. The top 5 sex-by-diagnosis interaction associated modules in MSSM-Penn-Pitt and NIMH-HBCC cohorts showed substantial overlap indicating reproducibility across cohorts (**Figure 5C**). Gene set enrichment analysis for the top 5 ranked modules identified 17 pathways that were common between MSSM-Penn-Pitt and NIMH-HBCC cohorts, including neural nucleus development, neuron projection morphogenesis, and regulation of neural precursor cell proliferation (**Figure 5D**).

Module H935 showed the most significant enrichment for sex-by-diagnosis differential expression signatures in the NIMH-HBCC cohort and it was nominally significant for association with schizophrenia common genetic risk variation (**Supplementary Table 2**). Moreover, its corresponding modules in the MSSM-Penn-Pitt cohort (C563 and C1055) were also ranked as the top enriched modules for sex-by-diagnosis differential expression signatures (**Figure 5C**). Module C1055 was one of the child modules within the module C563 (**Supplementary Table 2**). To better understand the sex-schizophrenia interaction network, we investigated the network topological structure of H935, comprised of 208 genes, and its MSSM-Penn-Pitt counterpart, C563, comprised of 105 genes (**Figure 6**). We identified 10 (*CEP170B, PRR12, CASKIN1, FBXO41, DLGAP4, AGAP2, NRXN2, NACC1, PACSIN1* and *PRRC2A*) and 7 (*TNRC18, LTBP3, ADGRB1, CASKIN1, ZDHHC8, APC2* and *PLEC*) key regulators of H935 and C563, respectively, using multiscale hub analysis in MEGENA. *CASKIN1* is the key regulator in both H935 and C563 and is involved in signal transduction pathways as a synaptic scaffolding protein (37, 38).

**Figure 6.**
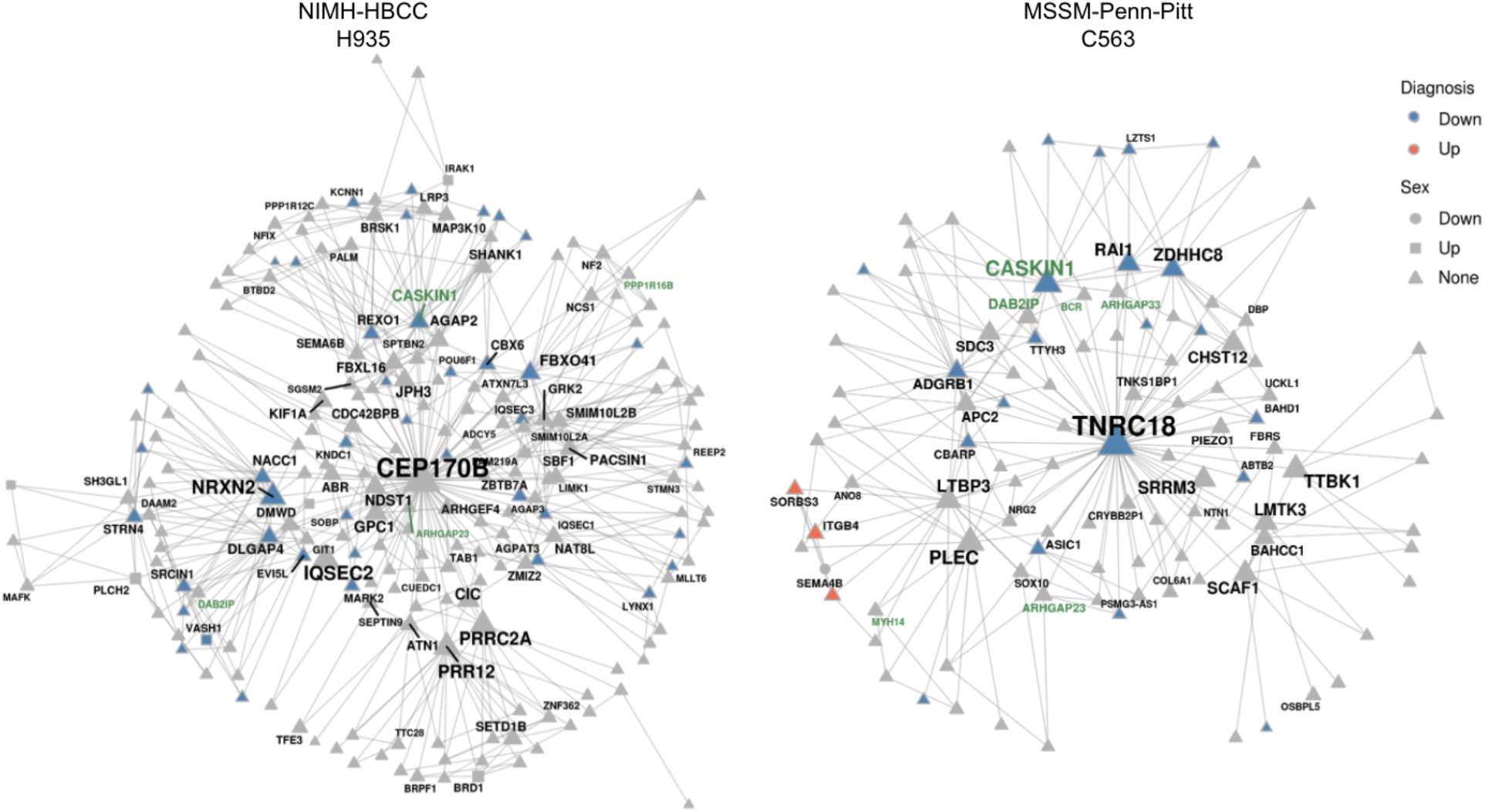
Co-expression modules enriched for sex-by-diagnosis differential expression signatures. The gene-gene interaction of the top-ranked modules H935 (left panel) and C563 (right panel) in two cohorts. Colors of nodes represent significantly up- and down-regulated genes in diagnosis analysis. Shapes of nodes represent dysregulated genes in sex analysis. Sizes of nodes represent the degrees of nodes (the number of connected nodes). Text labels in green represent the overlapping genes between two modules.

## Discussion

Despite the well characterized differences in onset and clinical presentation of schizophrenia between males and females, a mechanistic understanding of these differences has been lacking. To address this knowledge gap, we analyzed RNA-seq data from 437 controls and 341 cases from the CommonMind Consortium (3, 8) across four brain banks and two cohorts to determine differential expression signatures for diagnosis, sex, and sex-by-diagnosis interaction. Compared to the previous largest study that examined sex differences in schizophrenia (7), our cohort size increased by almost two folds. We first determined differential expression signatures for diagnosis. Meta-analysis across cohorts further improved the robustness of our findings. We found high concordance with schizophrenia signatures from other analyses (although we note that there is substantial overlap in samples between the two resources) and intermediate concordance with signatures for bipolar disorder and autism spectrum disorder (3, 4). Importantly, we find low similarity to signatures of Alzheimer’s’ disease, which affects similar brain regions, but has a vastly different disease mechanism.

We used these data resources to understand gene expression differences between males and females in the brain. Our analysis finds strong reproducibility across cohorts and meta-analysis identified a signature of 686 DEGs, including differential expression of 606 autosomal genes, which is widespread throughout the genome. In addition to dosage compensation and androgen signaling pathways, which were expected to show differences by sex, we also see enrichment for neurexin family protein binding and synaptic organization among sex differential expression. Stereological and microscopy studies have demonstrated that males have a significantly higher synaptic density than females in neocortex (39) and medial amygdala (40). Differences in the microanatomical and gene expression substrate of synaptic organization might contribute to the functional sex differences in brain activity. It is important to note that a simpler model, which does not include the cell-type composition, finds more significant DEGs (data not shown here). However these signatures were not reproducible across cohorts, presumably because they were affected by cell-type confounding. Thus, having two different cohorts provides a valuable resource to ensure the validity of our findings, but requires robust exploration and consideration of additional covariates (notably cell type composition) that might introduce artifactual cohort-specific effects.

Recent work by the GTEx Consortium identified 1,943 genes that are differentially expressed between males and females in the prefrontal cortex [BA9] after controlling for cell type composition (41). Despite assaying only 48 females and 127 males in this tissue, Oliva et al (41) used a sophisticated statistical method (42) to borrow information across 44 tissues from 838 individuals in the study, and identified significant genes using the local false sign rate (43) to dramatically boost statistical power. Our sex-specific findings show high concordance with the GTEx analysis (**Supplementary Figure 6**). Considering our combined results after meta-analysis as the discovery cohort, the replication rate in GTEx is estimated to be 96.1% (95.5% for autosomal genes). Considering GTEx BA9 as the discovery cohort, the replication rate in our dataset is estimated to be 80.7% (79.6% for autosomal genes) (see **Supplementary Methods**).

Studies involving a large number of individuals can be challenging to characterize due to complicated biological regulations underlying complex diseases. We did not find a genome-wide significant sex-by-diagnosis signature after multiple testing corrections, indicating that the effect size is small relative to the separate effect of diagnosis and sex. On the other hand, we provide two outcomes that support that sex-by-diagnosis signatures affect human brain transcriptome. First, there was a small, but significant correlation between interaction test statistics across the two cohorts. Second, by performing a network analysis to reduce dimensionality and elucidate interactions among genes, we found an enrichment for sex-by-diagnosis differential expression signatures that was highly reproducible across the two cohorts. The gene modules that were most associated with sex-by-diagnosis signatures involved a number of diverse pathways, including neural nucleus development, neuron projection morphogenesis, and regulation of neural precursor cell proliferation. For the top sex-by-diagnosis interaction enriched module, we identified a significant enrichment for schizophrenia common genetic variation. *CASKIN1* is a key regulator of this module that is reproduced among both cohorts. *CASKIN1* has a reduced expression in the schizophrenia patients, and is also known as a synaptic scaffolding protein to play a role in signal transduction.

Lack of genome-wide sex-specific findings should not be interpreted as the molecular etiology of schizophrenia in males and females is identical. Rather, the results indicate that any sex differences in disease signature are likely small and will require additional analyses in larger sample sizes. This notion is further supported by recent work on large-scale genome-wide association studies (GWAS) of schizophrenia (44–46). These studies did not identify genome-wide significant effect size differences between males and females and the genetic correlation between males and females is statistically not different from 1.0. More broadly, in analyses of large-scale GWAS and biobank data, findings of sex-specific effects or differences in heritability between males and females have been limited (44, 46, 47). These negative results from otherwise well-powered datasets underscore the challenge of studying the molecular mechanisms underlying clinical differences in schizophrenia between males and females.

The current analysis of bulk post mortem RNA-seq data from 497 males and 281 females has a number of limitations. Chiefly, our analysis implies that a sex-by-diagnosis interaction effect exists, but we are underpowered to detect it genome-wide with the current sample size. Additionally, ascertainment bias in post mortem studies means that far more male than female individuals are represented, which reduces the effective sample size and statistical power. Adding further data from resources generated from the PsychENCODE Consortium and other large-scale efforts, as well as specifically sampling more female samples, should be undertaken in the future to improve the power to detect sex-specific effects of disease. Furthermore, analysis of bulk tissue is necessarily limited to identifying effects that are either shared across cell types or large in a common cell type. It may be that sex-specific effects are concentrated in specific cell types. Ongoing work by PsychENCODE Consortium and others to generate large-scale single nucleus RNA-seq data from post mortem brains will have more power to identify cell type specific effects. Moreover, incorporating additional data types and analysis methods to integrate gene expression with genetics to identify sex-specific regulatory effects (41) may help to better understand sex differences in disease etiology.

In conclusion, the current study is the largest gene expression analysis in the human brain exploring sex differences in schizophrenia. Our results indicate that the effect size of sex differences in schizophrenia gene expression signatures is small. This further underscores the challenge of identifying robust sex-by-diagnosis signatures, which will require future analyses in larger cohorts.

## Acknowledgements

We thank the patients and families who donated material for these studies. This work was supported by the National Institutes of Health (R01AG050986 Roussos, R01MH109677 Roussos, R01MH109897 Roussos, U01MH116442 Roussos/Dracheva, R01MH110921 Roussos/Chess) and the Veterans Affairs (Merit grant BX002395 Roussos). J.B. was supported in part by NARSAD Young Investigator Grant 27209 from the Brain & Behavior Research Foundation. G.E.H. was supported in part by NARSAD Young Investigator Grant 26313 from the Brain & Behavior Research Foundation. Further, this work was supported in part through the computational resources and staff expertise provided by Scientific Computing at the Icahn School of Medicine at Mount Sinai and the assistance of members at the Mount Sinai Flow Cytometry CoRE. The funders had no role in the design and conduct of the study; collection, management, analysis, and interpretation of the data; preparation, review, or approval of the manuscript; and decision to submit the manuscript for publication. Human brain tissues provided by HBCC were supported by NIMH-IRP funding ZIC MH002903-14.

The authors declare no competing interests.

## Contributions

Designed study: G.E.H., M.P., S.D., J.F.F., A.C., B.D., S.K.S., P.R.

Performed data analyses: G.E.H., Y.M., K.S.M., J.B. Supervised data analysis: G.E.H., B.D., S.K.S., P.R. Data organization: M.A.P., K.S.M.

Wrote and edited the manuscript: G.E.H., Y.M., S.K.S., P.R.

Prepared and generated data from the samples: M.K.J., A.K. J.F.F, S.D., A.C, P.R.

## Supplementary Figures

**Supplementary Figure 1.**
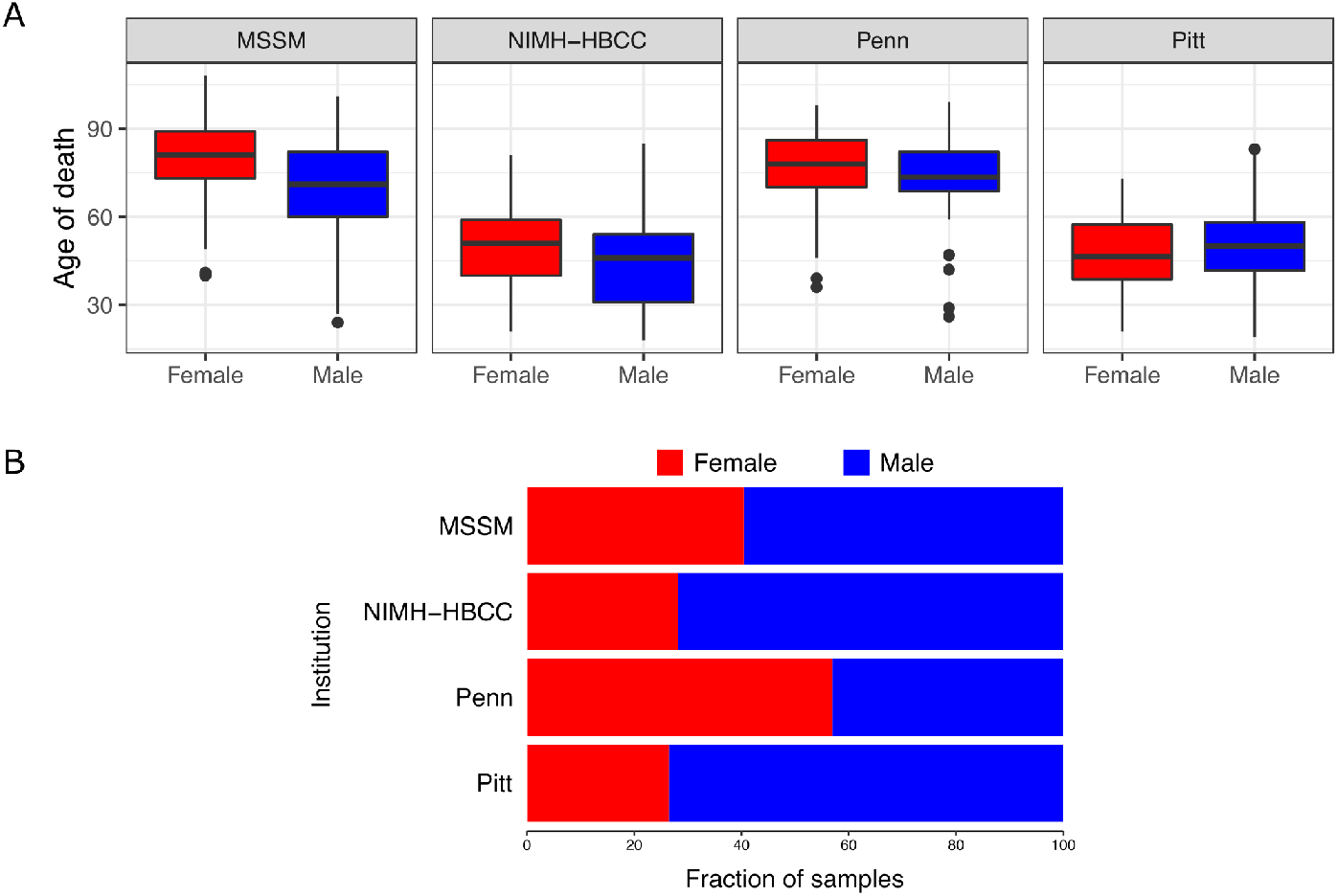
Illustration of confounding between sex, age of death and institution. **A)** Boxplot showing age distribution of males and females across the four institutions. Females tend to be older across most cohorts. **B)** Barplot showing the fraction of males and females across the four institutions. Females are underrepresented, especially in Pitt and NIMB-HBCC brain banks.

**Supplementary Figure 2.**
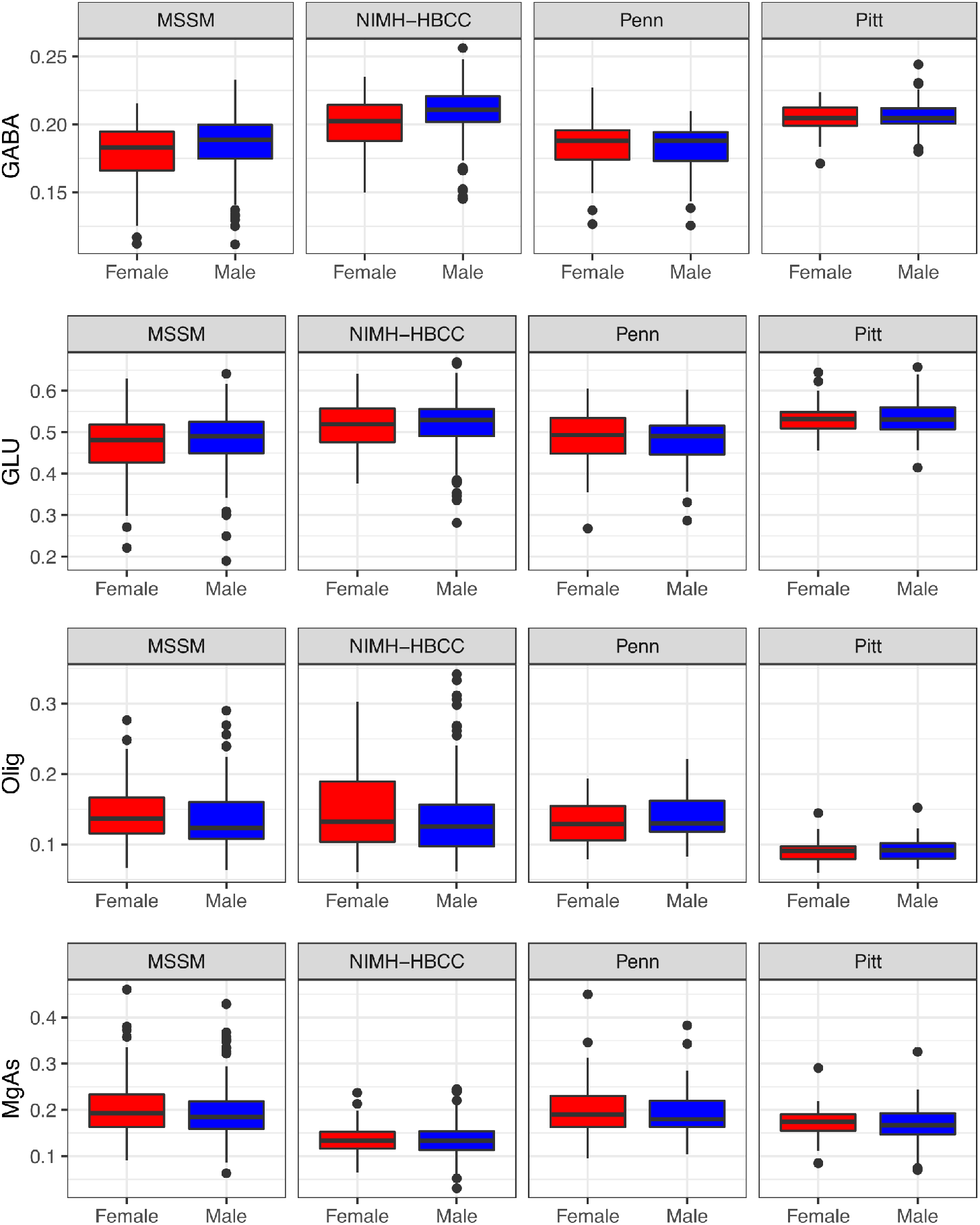
Illustration of confounding between sex, and estimated cell fraction. Females have a lower estimated fraction of GABAergic neurons, especially in MSSM and NIMH-HBCC brain banks.

**Supplementary Figure 3.**
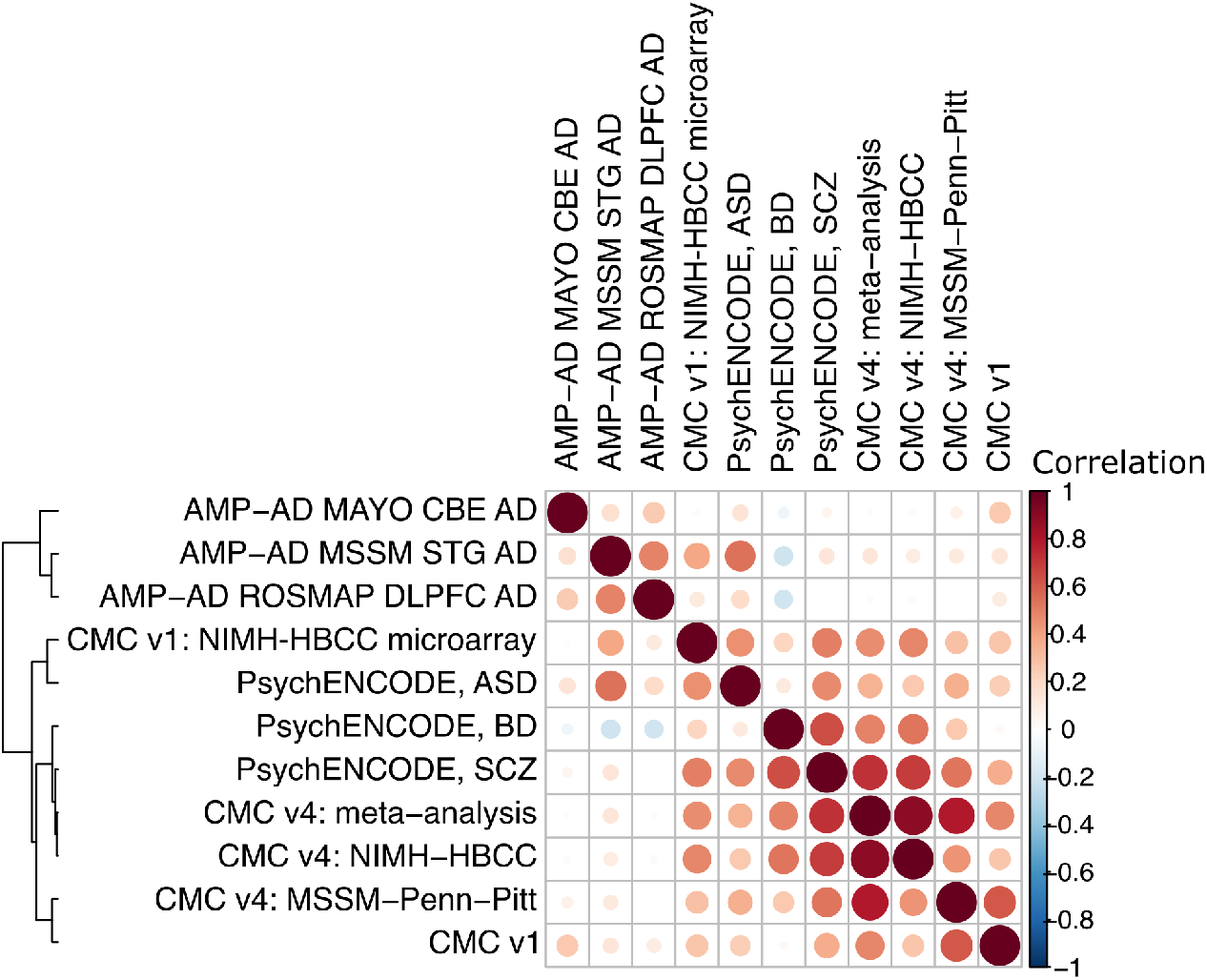
Concordance of disease signatures from post mortem brains. Correlation matrix and clustering of t-statistics from differential expression signatures from the current work compared to previously published disease signatures from post mortem brains.

**Supplementary Figure 4.**
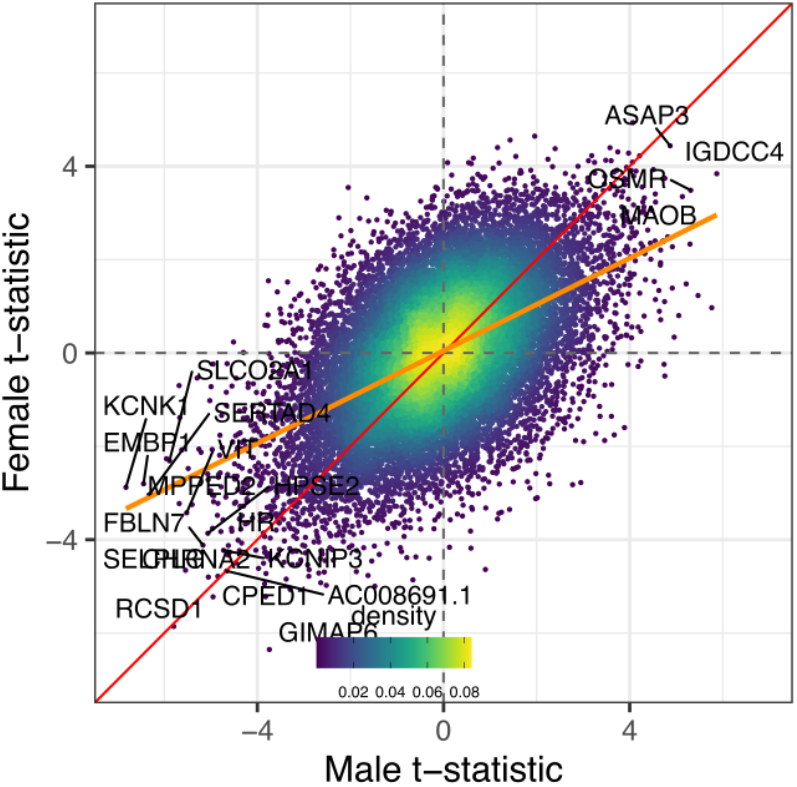
Concordance analysis between t-statistics from males and females when analyzed separately. Orange line indicates best fit from linear regression. Color of points indicates the density in the local region.

**Supplementary Figure 5.**
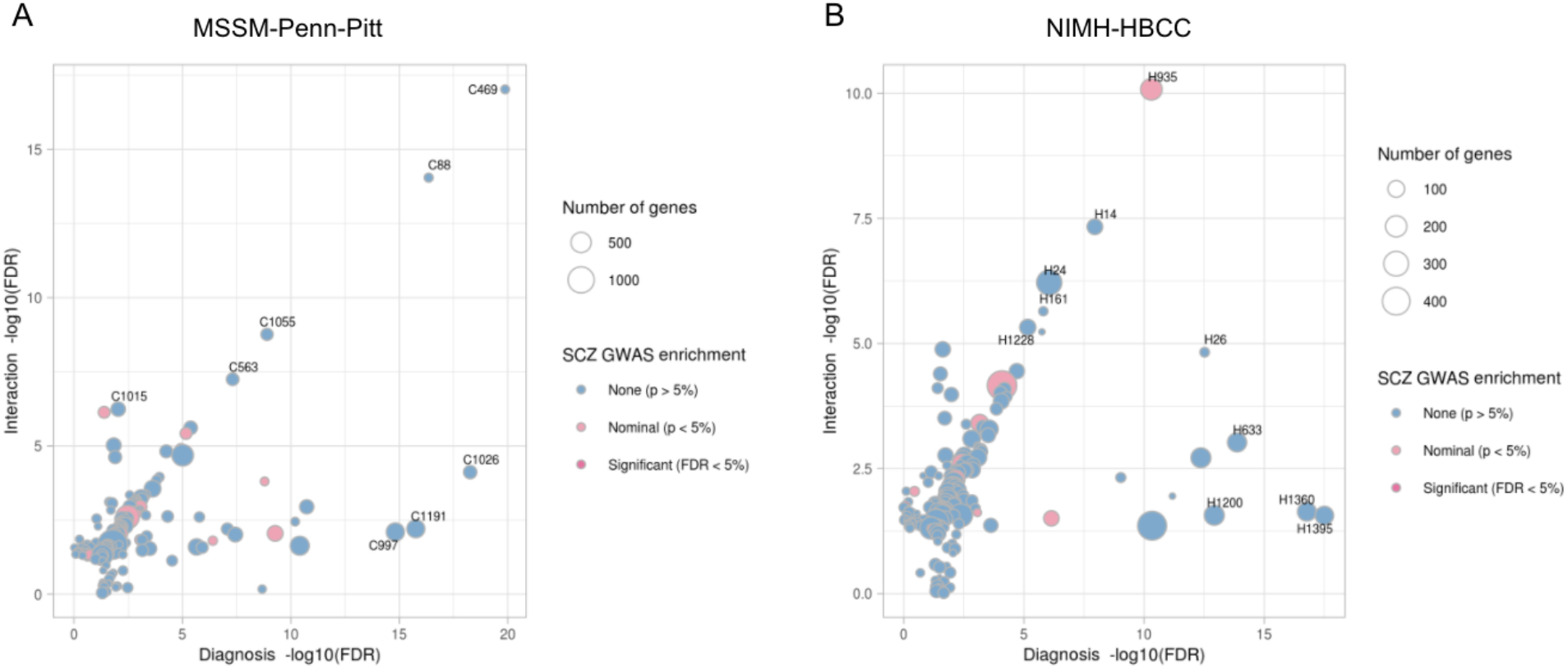
Concordance analysis showing FDR for each module enriched with diagnosis and interaction differential expression signatures in **A)** MSSM-Penn-Pitt and **B)** NIMH-HBCC cohorts. The top 5 enriched modules of diagnosis and interaction were labeled. Colors indicate the significance of modules enriched with schizophrenia (SCZ) GWAS. Blue represents modules not significantly enriched with SCZ; light pink represents modules enriched with SCZ at nominal p-value < 5%; dark pink represents modules significantly enriched with SCZ at FDR < 5%. The size of the nodes represents the number of genes within modules.

**Supplementary Figure 6.**
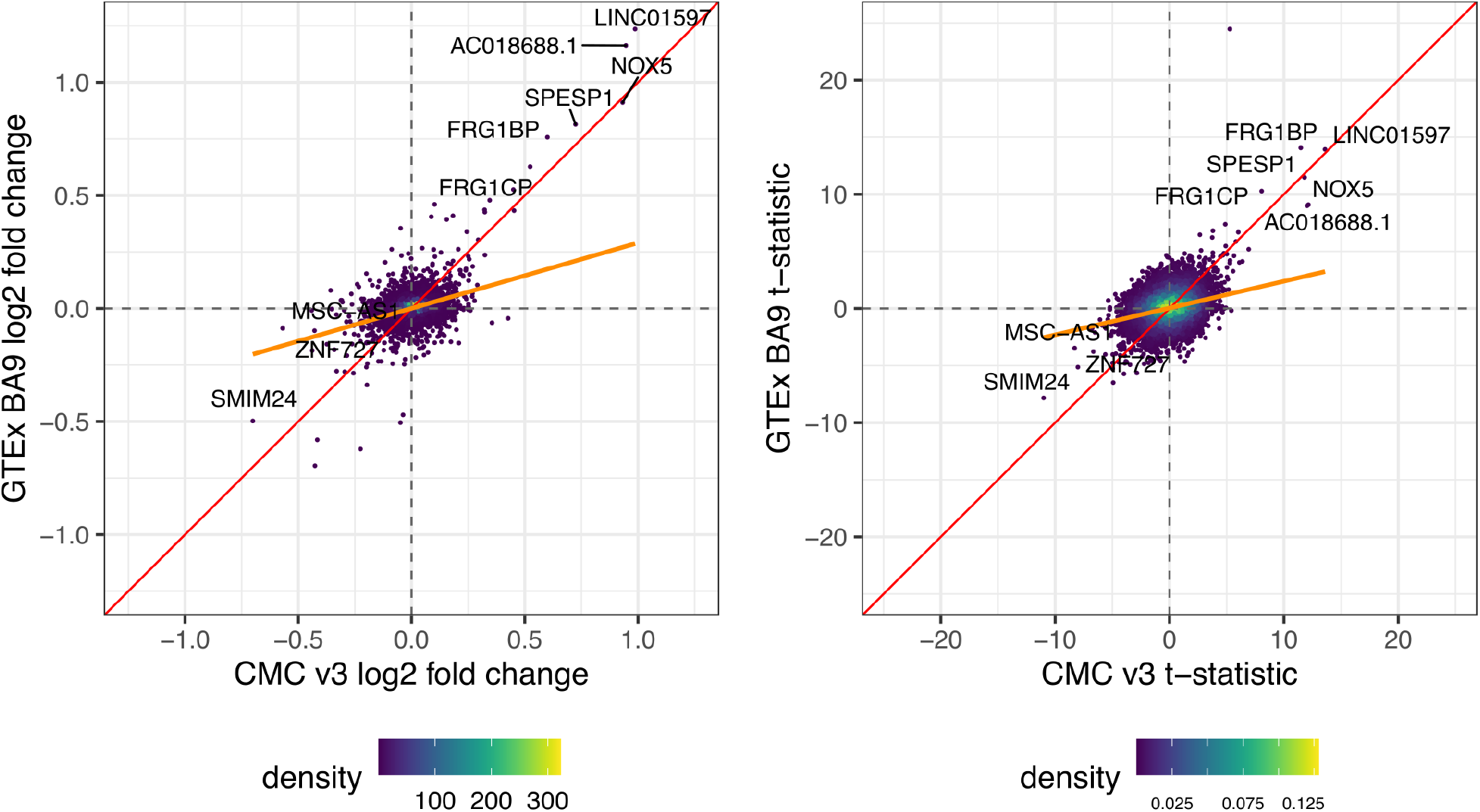
Concordance of differential expression between sexes between current dataset and frontal cortex [BA9] (BRNCTXB) from GTEx. **A)** Concordance analysis showing log_2_ fold change for each autosomal gene from CMC v4 and GTEx cohorts. We note that log_2_ fold changes from GTEx are the posterior estimates using a sophisticated method that borrowed information across 44 tissues. Orange line indicates best fit from linear regression. Color of points indicates the density in the local region. The genome-wide Spearman correlation between t-statistics is 0.223 (p < 2.35 x 10^−201^) and 0.218 (p < 1.59 x 10^−187^) for autosomal genes. **B)** Concordance analysis showing t-statistics for autosomal genes. For GTEx t-statistics here were generated using the posterior estimates of the log_2_ fold change and standard error from a joint analysis of all tissues.

## Supplementary Tables

**Supplementary Table 1.** Number of RNA-seq samples. Counts are shown for both cohorts per institution and are stratified by case/control status and sex.

| Cohort | Institution | Female |  | Male |  |
| --- | --- | --- | --- | --- | --- |
|  |  | Controls | SCZ | Controls | SCZ |
| MSSM-Penn-Pitt | MSSM | 77 | 46 | 84 | 97 |
|  | Penn | 19 | 34 | 18 | 22 |
|  | UPitt | 23 | 13 | 59 | 41 |
| NIMH-HBCC | NIMH | 40 | 29 | 117 | 59 |
| **Total** |  | **159** | **122** | **278** | **219** |

**Supplementary Table 2. Modules associated with the differential expression signatures and the schizophrenia GWAS.**

https://docs.google.com/spreadsheets/d/1f_SPRysprG5yZc9jirxRwNM7tDRFe67dKlF-7Ny2hRQ/edit?usp=sharing

## Supplementary Methods

### Estimating replication rate using GTEx results

The π_1_ statistic is widely used to estimate the replication rate. The approach works by identifying a set of tests where the null is rejected in the discovery dataset, and using frequentist p-values in the replication cohort to estimate the fraction of tests where the null hypothesis is rejected. Thus p-values are required in order to apply this method.

However, the GTEx analysis (41) used an empirical Bayes approach called MASH (42) to borrow information across all 44 tissues. For every gene and tissue, MASH reports posterior estimates of the log_2_ fold change and its standard error. Significant genes are called using the local false sign rate (lfsr) which indicates the posterior probability that the sign of the estimated log_2_ fold change is wrong (42, 43). Since lfsr values, rather than p-values, are used to evaluate each hypothesis of differential expression, an alternative method must be used to evaluate the replication rate.

Here we propose an approach very similar to the π_1_ statistic, except that the fraction of tests estimated to reject the null hypothesis is computed from the set of lfsr values. Let *S* be the set of tests called significant in the discovery cohort and let *lfsr_i_* be the lfsr value for test *i* in the replication cohort. Since lfsr values are posterior probabilities, ∑_*iϵs*_ *lfsri* is the total probability mass over the *k* tests supporting the null hypothesis. Similarly, ∑_*iϵs*_(1 − *lfsr_i_*) is the total probability supporting the alternative hypothesis so that 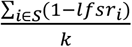 can be interpreted as the fraction of null hypotheses that are rejected.

In order to be consistent in our comparison of GTEx and CMC v4, we used the ashr package (43) to generate lfsr values for CMC v4 sex signatures.

